# DNA methylation contributes to plant acclimation to naturally fluctuating light

**DOI:** 10.1101/2024.06.07.597890

**Authors:** Robyn A Emmerson, Ulrike Bechtold, Nicolae Radu Zabet, Tracy Lawson

## Abstract

Plants in the natural environment experience continuous dynamic changes in light intensity. We have limited understanding on how plants adapt to such variable conditions. Here, we exposed *Arabidopsis thaliana* plants to naturally fluctuating light regimes alongside traditional square light regimes such as those often found in control environment growth chambers. The physiological response was highly consistent across experiments, indicating the involvement of an epigenetic mechanism, leading us to investigated differences in DNA methylation. Our results identified a large number of alterations in DNA methylation patterns between fluctuating light acclimated plants, and square light acclimated plants, demonstrating natural fluctuations in light impacts the plant epigenetic mechanisms. Most importantly, there are more differences in DNA methylation patterns between different light pattern regimes than between different light intensities. These differences in DNA methylation were accompanied by significant changes in gene expression, some of which correlated with altered DNA methylation. Interestingly, several transposable elements which displayed differential methylation were found to be differentially expressed between light regimes. Our data suggests DNA methylation plays a role in acclimation to natural light which may directly regulate gene expression and impact transposable element activation.

## Introduction

In natural environments, plants are continually exposed to changing environmental conditions, including rapidly altering light intensity due to weather and shading from overlapping leaves, as well as diurnal and seasonal effects (1–3). Light is crucial for photosynthesis and variations in its quantity, quality and timing significantly impact on plant growth, development and productivity (4–6). To manage these potentially stressful fluctuations, plants undergo acclimation, a process involving alterations in molecular physiology. Although, numerous studies have examined the impact of light intensity, the mechanisms for acclimation to dynamic irradiance have received limited attention to date.

Acclimatory responses have been correlated with epigenetic changes as both a response to the stress (7–10) or as a priming effect (11, 12). Transgenerational epigenetic inheritance in plants is well documented and can provide a mechanism for this priming effect. In particular, maintenance of CpG methylation by MET1 and DDM1 is essential for transgenerational epigenetic inheritance (13–16). The dynamic nature of DNA methylation, and its potential to control gene expression, has previously been studied in plant stress responses and acclimation (17, 18) as it represents a fast and changeable mechanism by which plants can respond to their environments.

Nevertheless, there is currently limited evidence for the role of DNA methylation in light stress responses and acclimation. Short-term light stress, with a small number of fluctuations in light, have been reported to have limited effects on DNA methylation (19). However, production of reactive oxygen species (ROS), including hydrogen peroxide (H_2_O_2_), as a response to high light has been linked to alterations to the methylome. In tobacco mutants, which overproduce H_2_O_2_, loss of DNA methylation was observed in comparison to the control plants (20), suggesting overproduction of H_2_O_2_ correlates with loss of methylation.

More generally, ROS have been implicated in epigenetic reprogramming. For example, MutS Homologue 1 (MSH1) present in sensory chloroplasts has been implicated in alterations to DNA methylation. Knockout of MSH1 results in genome-wide reprogramming of DNA methylation (21), further suggesting a role for chloroplast signalling in epigenetic change. Combined with knowledge that high light stress results in ROS production (22, 23), this indicates that light environments can impact the plant methylome. There is also evidence that small RNAs (sRNAs) are induced in response to high light in *Arabidopsis* and impact gene expression (24). Since 24nt sRNAs are capable of guiding DNA methylation via the RdDM pathway (25), this suggests another possible mechanism by which the light environment can impact DNA methylation.

Although these studies investigate excess light stress or peaks and troughs in light, none are reflective of environmental light regimes where the fluctuating frequency is much greater. Our previous work showed that the physiology of *Arabidopsis* is impacted by a naturally fluctuating light regime (26, 27), which indicates a possible involvement for DNA methylation. Here we have utilised naturally fluctuating light regimes (27) to assess the impact of acclimation on DNA methylation and gene expression in *Arabidopsis thaliana* (Fig.1). Our results showed large differences in DNA methylation patterns of plants exposed to fluctuating light regimes compared to plants exposed to square wave light. In particular, we found hundreds of differentially methylated regions in CpG and non-CpG context and these regions consist of both genes and transposable elements. Furthermore, we linked these epigenetic changes to changes in transcription that were able to explain the observed phenotypes.

**Fig. 1:**
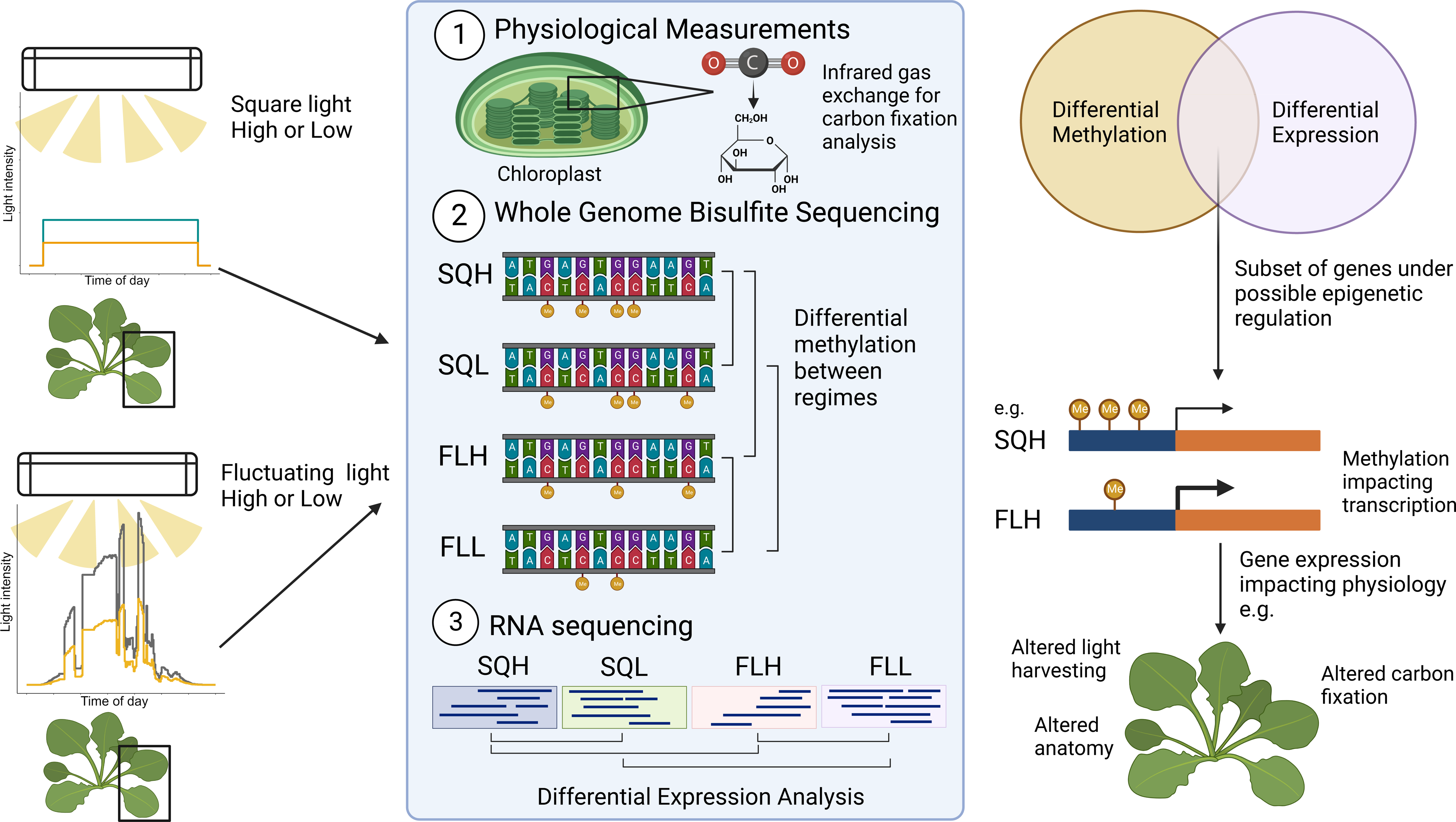
Schematic representation of experimental design used in this study. Arabidopsis thaliana. (Col-0) were exposed to one of four light regimes, two representing laboratory light conditions of a high and low intensity (SQH and SQL), and two representing natural light conditions (FLH and FLL). These were then subjected to physiological assessment, followed by DNA methylation (WGBS-seq) and transcriptome (RNA-seq) analysis to determine whether light regime impacted epigenetic landscape, and whether there was a correlation between DNA methylation and gene expression which could be associated with the altered physiology.

## Results

### Growth light regime induces loss of DNA methylation

To investigate the impact of growth light regime on DNA methylation, plants were acclimated to square and fluctuating light regimes. In both cases, we considered high and low light, which resulted in four conditions: square light high (SQH), square light low (SQL), fluctuating light high (FLH) and fluctuating light low (FLL) (Fig. 1). The plants were then subjected to physiological assessment (Fig. S1) to ensure a consistent phenotype with previous data (27). Our results showed that there is little change to the global methylation profiles between the four growth conditions (Fig. 2A and Supplementary Fig. S2), which indicates that light acclimation is not resulting in global changes to the epigenetic machinery, but instead may be impacting DNA methylation at specific regions in the genome.

**Fig. 2:**
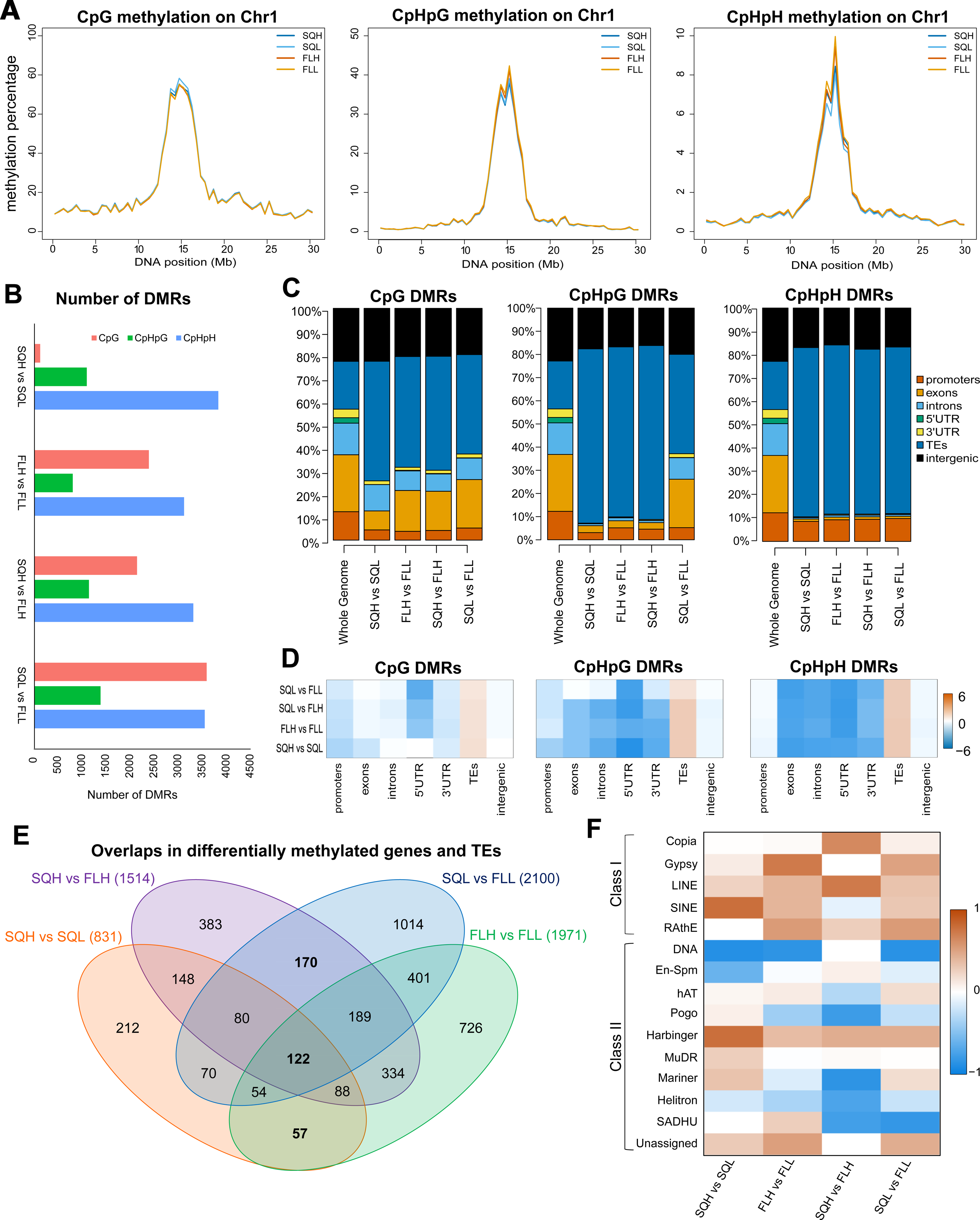
The effects of light regime on DNA methylation. *(A)* Low resolution profiles of DNA methylation in the four light conditions. We considered separately CG, CHG and CHH methylation patterns and only plot methylation data on chromosome 1. *(B)* Number of DMRs between the different conditions. *(C)* Annotation of the DMRs to different genomics features (promoters, exons, introns, 5’UTRs, 3’UTRs, TEs and intergenic regions) and *(D)* enrichment computed as log_2_(observed/expected) for the genomic features. *(E)* Venn diagram of the overlaps of the genes and TEs that display differential methylation between the four light regimes. *(F)* TEs subfamilies and evaluation of the change in DNA methylation. TEs that gain methylation (1) are marked by red and TEs that lose methylation (−1) are marked by blue.

Next, we computed the Differentially Methylated Regions (DMRs) between square and fluctuating light regimes and identified between 119 and 3,574 DMRs in CpG, 793 and 1,373 DMRs in CpHpG and 3,105 and 3,814 DMRs in CpHpH contexts (Fig. 2B and Fig. S3). This is significantly higher compared to previous reports where plants were acclimated for 7-8 days (19), which indicates that the level of epigenetic changes might be impacted by the duration of the acclimation to fluctuating light regime. Interestingly, we found that more epigenetic changes are observed between square light and fluctuating light regimes than between low and high light. This suggests that plants display a stronger acclimation when changing the light frequency/pattern than to changes in the amplitude of the light. Some of these regions gain DNA methylation, while other lose it, but there is no clear trend for that in the four comparisons we performed (Fig. S3).

Functional annotation of these DMRs indicates that acclimation could be linked to epigenetic changes in transposable elements and intergenic regions (Fig. 2C-D). In some specific cases, we also found a high number of DMRs in introns and promoters. These together with intergenic regions are potential regulatory regions, indicating they are involved in gene regulation. Transposable Elements (TEs) accounted for most DMRs across all contexts, indicating TE regulation is an important mechanism during acclimation to all growth light regimes. Furthermore, if we consider all three methylation contexts, DMRs are only enriched at TEs (Fig. 2D).

Together, our data suggest that DNA methylation profiles changed in response to different growth light regimes, with differences between the light frequency resulting in a higher number of DMRs compared to differences in light intensity.

### Exposure to fluctuating light regimes leads to hypermethylation of transposable elements

Next, we investigated if there are common changes in DNA methylation at genes and TEs occurring between the four different light regimes (Fig. 2E). Our results showed that most of the DMRs within genes or TEs were unique to the light regime, suggesting there is a distinct epigenetic change associated with acclimation to the growth light regimes. The highest number of unique DMRs were noted in SQL vs FLL, suggesting a greater degree of regulation may be required under fluctuating low light conditions. 122 genes and TEs (called light constitutive genes) were differentially methylated across all regimes, indicating these may be essential to growth light acclimation, regardless of the actual light regime (Fig. 2E and Table S1). Furthermore, 170 differentially methylated genes and TEs (called light frequency genes) were shared between square and fluctuating regimes independent of the light intensity, but were not identified between high and low light regimes. Finally, 57 differentially methylated genes and TEs (called light intensity genes) were shared between high vs low regimes independent of the light frequency, but not between square and fluctuating regimes. Of the 122 light constitutive genes, the majority (90) were TEs (Table S1), meaning TE regulation could have a role in the acclimation response. Gene functions included Vacuolar H+ ATPase (AT3G58730), as well as proteins associated with signalling in the chloroplast (AT1G19090) and leaf senescence (AT1G54040). When the overrepresentation analysis of these groups was considered, no functional groups were found to be significantly enriched (FDR<0.05).

Fig. 2C showed that a large proportion of DMRs between different light regimes overlapped TEs. To further investigate this, we considered different types of TEs and found that DMRs were enriched in the majority of retrotransposons (Class I TEs) and depleted in DNA transposons (Class II) (Fig. 2F). For example, LINE TEs were enriched across all regimes, while Copia were only enriched in DMRs between square and fluctuating light, independent of the light intensity. Gypsy and SINE TEs were enriched in all cases except between square high and fluctuating light high. This indicates that acclimation to light can result in changes in DNA methylation at specific classes of retroelements.

### Epigenetic changes upon exposure to light regimes correspond to activation in transcription

To determine whether the DNA methylation is accompanied by transcriptional changes in the plants acclimated to fluctuating light, we performed RNA-seq in three biological replicates for the four light regimes. Principal component analysis (PCA) analysis confirmed that the biological replicates grouped together (Fig. 3A). The groups of FLH and FLL clusters displayed a greater degree of spatial separation in comparison to SQH and SQL, indicating that gene expression displays higher variability in fluctuating light compared to square light regimes, even in the case when the fluctuating light pattern is the same for all three biological replicates (also see Fig. S4).

**Fig. 3:**
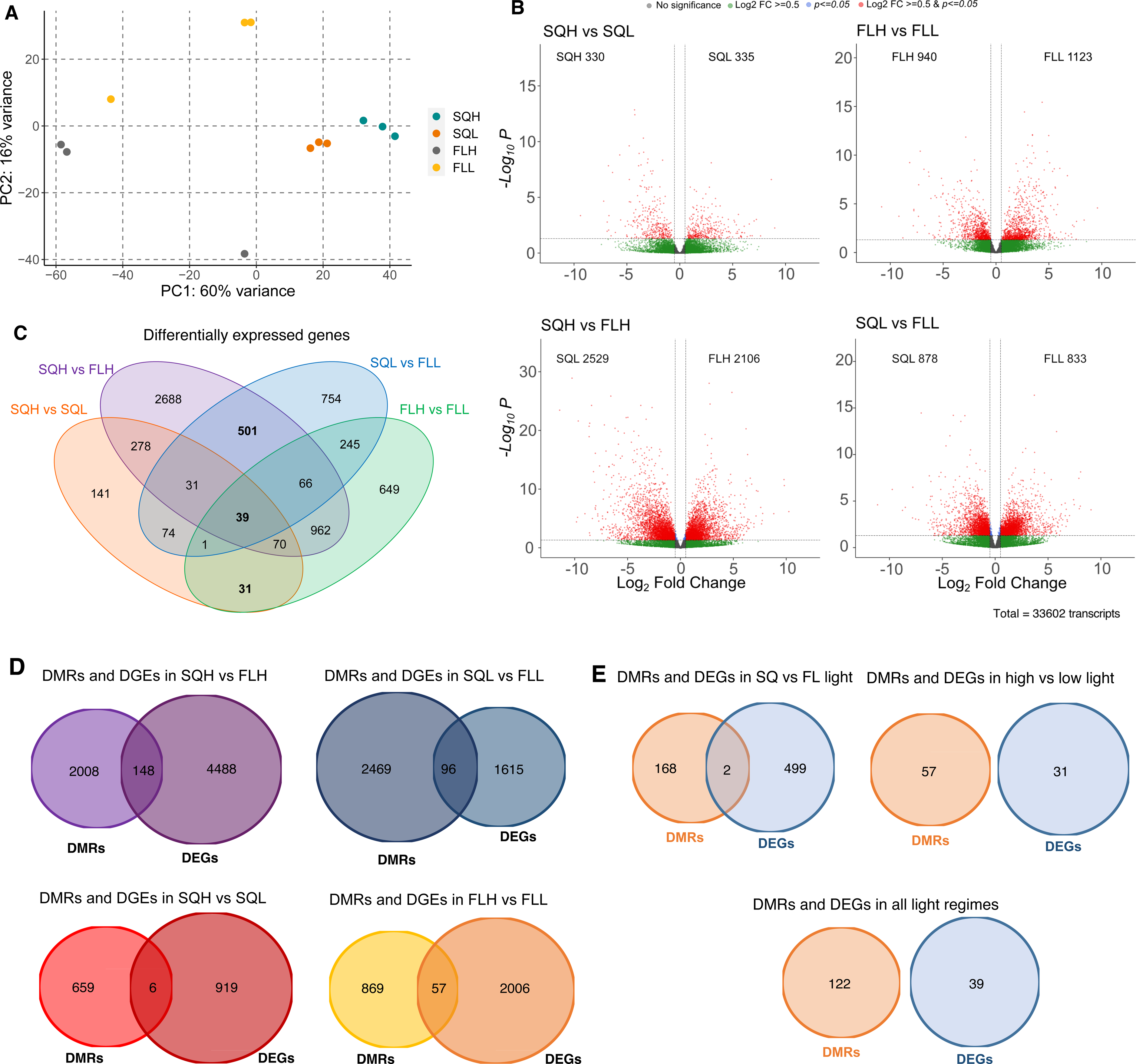
Transcriptome of plants exposed to the different light regimes. *(A)* PCA analysis of RNA-seq data in three biological replicates for the four light conditions. *(B)* Volcano plots for the significantly differentially expressed genes (DEGs). The plot highlights the genes that show differential expression between the two light regimes in the header using a p-value threshold of 0.5 and a log_2_ fold change threshold of 0.5. The number in the corresponding inset represents the number of genes with higher expression in the corresponding light regime. *(C)* Venn diagram identifying common and specific set of genes differentially expressed in the four comparisons. 501 genes are differentially expressed in fluctuating compared to square light regimes, 31 in high compared to low light regions and 39 between all comparisons. *(D)* Venn diagrams of genes that are differentially expressed in one of the four comparisons and also overlap with a DMR (in any context) in the corresponding comparison. *(E)* Genes that are differentially expressed and overlap with DMRs in the three combined comparisons, namely: *(i)* fluctuating vs square light, *(ii)* high vs low intensity and *(iii)* all comparisons.

Next, we investigated if genes are differentially expressed between the four light regimes and found that between hundred and thousands of genes are indeed differentially expressed (DEGs) (Fig. 3B). The lowest number of differentially expressed genes (665) were identified between SQH and SQL, indicating that changing light intensity upon exposure of a square light leads to lowest number of transcriptional changes. In contrast, comparing SQH to FLH lead to highest number of differentially expressed genes (4,635), which means that changes in light frequency had the largest effect on the plants. Overall, our results indicate that acclimation to fluctuating light requires a greater magnitude of change than acclimation to light intensity. To assess whether a subset of these genes is shared across regimes, differentially expressed genes were grouped depending on regime (Fig. 3C). 39 genes were found to be differentially expressed across all regime comparisons, indicating these genes may be key for light acclimation. A further 31 genes were shared between high and low light intensity (SQH vs SQL and FLH vs FLL), and 501 genes were differentially expressed between square light and fluctuating light (SQH vs FLH and SQL vs FLL). Gene ontology analysis revealed significant enrichment across 9 terms in the 39 genes differentially expressed in all regimes (Fig. S5A) most of which were associated with stress responses, suggesting light regime significantly impacts how the plants respond to stress. Nevertheless, only response to wounding term was enriched when comparing HL v LL (Fig. S5B), which suggests that higher light intensity could potentially induce a response similar to wounding without necessarily the physical wounds. Interestingly, when comparing SQ v FL conditions, we did not identify any stress response terms significantly enriched. However, metabolic and cell cycle terms were enriched when comparing SQ v FL (Fig. S5C), suggesting metabolism and meiosis are primarily impacted in plants acclimated to light patterns.

DNA methylation is linked directly to transcriptional repression or gene silencing (28, 29), but this is not always the case and it is now known that not all changes in DNA methylation translate into changes in gene expression (30–32). In plants, we know that there is transgenerational inheritance of DNA methylation (33, 34), which suggests the possibility that some of these transcriptomic changes can be epigenetically encoded and have the potential to be maintained in several generations. To investigate if the transcriptomic changes are linked with epigenetic state, we looked at genes that showed both differential expression and differential methylation in the different light regimes (Fig. 3D-E). Our results showed that there was a greater number of genes displaying both differential expression and differential methylation under SQH vs FLH and SQL vs FLL than under SQH vs SQL and FLH vs FLL (Fig. 3D). However, none of these were found to be significantly over-represented (Chi-squared test, p-value > 0.9). The genes displaying differential expression that are under possible control of DNA methylation exhibit wide functionalities (Table S2), with many related to plastid activity and stress responses as well as photosynthesis-related, demonstrating that light can have impacts on plant gene expression outside of photosynthesis.

When square light and fluctuating regimes were grouped together, 2 genes were found to be both differentially methylated and expressed (Fig. 3E), suggesting these may have a key role in fluctuating light acclimation. One was AT1G03090 (MCCA), a subunit of 3-methylcrotonyl-CoA carboxylase (MCCase) which catalyses a key step in the catabolism of leucine and isovaleric acid in the mitochondria (35, 36). Accumulation of MCCA is known to occur in response to darkness, while exogenous sucrose has been shown to decrease MCCA expression (37), suggesting there could be transcriptional control as a result of both light and sugar signalling. The second gene is AT1G27880, a DEAD/DEAH box RNA helicase family protein. Although this gene is not characterized, the general family contains a range of RNA helicases with specific expression patterns in different developmental stages (38). Expression of several family genes has been associated with abiotic stress tolerance, and reported to be induced by glucose, ABA, and salt (39, 40).

### Changes in light regimes lead to activation of TEs

Demethylation of TEs can lead to their activation (41–46). To determine whether the differential methylation has led to activation or silencing of TEs, we identified the set of TEs that were both differentially methylated and expressed in the different light regimes (Table S3). We found only 23 TEs that were differentially methylated and differentially expressed across light regimes. Changes in frequency of light (SQH vs FLH and SQL vs FLL) accounted for the majority of these TEs, 12 and 7 respectively. Under SQH vs FLH, 10 of the 12 TEs showed increase in expression, suggesting increased TE activation under SQH light compared to FLH light (of which 4 displayed loss of DNA methylation; ATIS11A, ATGP3, ATLANTYS1, and VANDAL20) and 2 loss of expression (of which 1 displayed gain of methylation; TA11). In contrast, under SQL vs FLL, 6 of the 7 TEs displayed a decrease in expression (of which 3 displayed gain of DNA methylation; ATGP1, TA11, and a hAT-like TE), with FLL light showing an increase in transcription of these TEs, and 1 TE displayed an increase in expression (coupled with a loss of DNA methylation; ATRE2). To assess whether methylation, and therefore change in expression, of these TEs have the potential to be stably inherited, this list was compared against a list of stable epialleles associated with transposable element sequences (13). Across the list, only one TE was noted to have this potential, SADHU, which was differentially methylated and expressed under SQH vs FLH conditions.

TEs can also act as regulatory regions (47, 48). To assess the possible impacts of changes TE DNA methylation levels on nearby gene expression, we also investigated the expression of genes within 1 Kb of differentially methylated TEs (Table S4). Interestingly, we found a stronger link between changes of methylation at TEs and changes in gene expression at nearby genes, with SQH vs SQL showing a total of 29 DEGs within 1 Kb of a differentially methylated TE, while FLH vs FLL had 110. In the square vs fluctuating comparisons, SQH vs FLH displayed the greatest number of DEGs (258), while SQL vs FLL yielded 97 genes. Despite the number of genes differentially expressed, gene ontology analysis revealed there was no significant enrichment across terms.

### Loss of DNA methylation improves photosystem II efficiency under fluctuating light

One possibility is that changes in DNA methylation only correlate to changes in gene expression upon exposure to different light regimes and DNA methylation is not required in order to observe these physiological differences in plants. If that is the case, then these physiological changes in the plants upon exposure to different light regimes would still be present in a DNA methylation deficient mutant plants. To further investigate the link between DNA methylation and fluctuating light acclimation, we performed chlorophyll fluorescence imaging on *met1-1* knockdown plants following 7 days of acclimation to SQH or FLH regimes. *met1-1* plants display approximately 75% loss of CG methylation, specifically in gene bodies (49). This allows us to investigate which differences in physiological response upon different light regimes require DNA methylation (in CG context) to be established and maintained. Interestingly, we observed a difference between the wild type and *met1-1* plants under both regimes, with reduced methylation improving the photosystem II operating efficiency (Fig. S6A) when acclimated to FLH light, with the opposite impact under SQH, although this was not significant. We also found a difference between SQH and FLH WT plants that was not identified in the *met1-1* mutants (Fig. S6A and S6D). This appeared to be due to the maximum efficiency of photosystem II (*F_v_’/F_m_’)* rather than photochemical quenching (*F_q_’/F_v_’)*, with decreases noted in the mutant relative to the wildtype under SQH, and a small increase in FLH acclimated *met1-1* compared to wildtype (Fig. S6B-C). Overall, our results indicate that at least some of the changes in plant physiology (and, consequently, in gene expression) under square light conditions do require DNA methylation. We also found a potential trade-off between photosynthetic efficiency and acclimation under FLH conditions.

## Discussion

DNA methylation is highly dynamic and known to respond to environmental stimuli. There is limited evidence for the effects of long-term light stress and acclimation on the methylome, with previous studies concluding there is little impact of light regime on DNA methylation despite clear physiological phenotypes (19). However, the impacts of natural light regimes and acclimation over the lifespan of *Arabidopsis* has not been conducted. This, combined with the known negative effects of fluctuating light on carbon assimilation in both tropical and crop plants (50–52), and the highly consistent phenotype exhibited under the regimes used here (26, 27), indicated a role for epigenetics.

The peaks and troughs of the fluctuating light regimes are likely to act as cues for epigenetic change. For example, during the peaks of light in both FLH and FLL regimes, ROS are likely to be generated (53). ROS generation has previously been associated with epigenetic change, and a recent study has correlated increased ROS due to abiotic stress with hypomethylation (54). Troughs may act as a recovery period for the plants, where ROS production is reduced, and so could result in re-methylation of the loci demethylated during high ROS. However, low light could also be considered as stressful for plants, prolonging vegetative growth (55) and reducing carbon assimilation capabilities (27), so itself could cause hypomethylation.

There is increasing evidence for the impacts of light on plant epigenetic profiles with direct impacts on gene expression. For example the histone demethylase IBM1, which removes methyl groups from H3K9 and acts to reduce non-CpG methylation, has been found to impact anthocyanin biosynthesis in response to high light (56). Following 48h high light, IBM1 expression was induced in wild type *Arabidopsis,* acting to increase expression of SPA which positively regulate production of anthocyanin biosynthesis(56). This was attributed to both demethylation of H3K9 and to local decreases in non-CpG methylation of the SPA genes (56). While there was no noted change in expression in IBM1 our study between light regimes, this provides evidence for direct epigenetic mechanisms associated with light exposure. Furthermore, more condensed chromatin has been associated with growth under light compared to dark conditions in soybean, with regulation of genes associated with photomorphogenesis particularly upregulated (57). Together with our data, this suggests light conditions can specifically impact epigenetics and gene expression. Due to the unclear nature of the causality and correlation between DNA methylation and gene expression, it is difficult to conclude what the impact of light on methylation and gene expression truly are (32, 58). It may be that methylation here acts as a priming event, so offspring from these plants are better able to cope with the parental light regime. Priming and transgenerational effects are well known to occur in plants as a response to multiple stresses, including cold stress and herbivory (59–61). Therefore, transgenerational maintenance of DNA methylation may represent a competitive advantage for offspring to grow under these light conditions.

Relatively few differentially methylated genes were found to be differentially expressed (Table S2) in this study. This is consistent with previous studies into the effects of stress. For example, only 9 of 1,562 differentially methylated genes were differentially expression under prolonged cold treatment in *Brassica rapa* (62), while only 31 of over 5,000 DMRs had differential expression under iron deficiency in rice (63). Together, these studies demonstrate that the magnitude of differential methylation between treatments is often larger than the resulting differential expression, suggesting differential methylation is not always controlling expression. Similar findings were noted here, with only a small subset of DMRs and DEGs overlapping (Fig. 3D-E).

Interestingly, we observed change in methylation status and expression at a range of transposable elements (TEs). Activation of TEs has been seen under heat stress in *Arabidopsis*, with novel insertions reported (48, 64, 65). However, while activation of TEs is a possibility, it is more likely that TEs act as *cis*-regulatory regions to nearby genes. We found that genes within 1 Kb of a differentially methylated TE were differentially expressed in different light regimes. Increased methylation of TEs has previously been shown to impact neighbouring gene expression. For example, in *Arabidopsis*, silencing of a SINE TE neighbouring FWA, a locus associated with flowering, results in FWA silencing (66). Methylation has been noted to spread from TEs into neighbouring genes and the repressive chromatin state resulting from this has been proposed to also impact on nearby genes (67). It is also possible that loss of methylation leads to increased expression of TE-neighbouring genes due to the opening of the chromatin structure.

Our study demonstrates the importance of light regime (the pattern in which light changes) for plant growth and development. Comparisons of light patterns (square light to fluctuating light) resulted in a greater level of both epigenetic and transcriptomic change in genes and TEs, indicating substantial differences in the way plants respond to dynamic light environments than to the intensity of the light (high compared to low). This could have significant impacts on studies aimed at field crops grown under laboratory conditions, as this provides evidence as to how and why plants may react differently in laboratory and field studies. Furthermore, our results indicate that acclimation to light patterns involves a greater level of epigenetic and transcriptomic changes that acclimation to light intensity, which is of relevance to field plants and current climate changes.

## Materials and Methods

### Experimental model and growth conditions

*Arabidopsis thaliana* Col-0 background were germinated in 5 cm^2^ pots on a peat-based compost (Levingtons F2S, Everris, Ipswich, UK), and placed in a controlled growth environment at 22°C, 65% relative humidity, 8/16h light/dark cycle, CO_2_ concentration 400 μmol mol^-1^. At 14-days, the seedlings were transplanted to individual 5cm^3^ pots containing the same soil as above and returned to the controlled environment.

At the 4-leaf stage, plants were removed from the controlled environment and placed under Heliospectra LED light source (Heliospectra, Göteborg, Sweden) programmed to each light regime (Supplementary Fig. 1) in a dark room maintained at 21°C/16°C Day/night, 50% relative humidity. Average light intensity for high light conditions was 460 μmol m^-2^ s^-1^ and 230 μmol m^-2^ s-^1^ for low light conditions on a 12h/12h day/night cycle. Plants were kept in well-watered conditions, with their position under the light source randomised every 3 days to remove any potential heterogeneity in spectral quantity and quality.

### Physiological Measurements

After 20 d acclimating to the light regimes, plants were subjected to infrared gas exchange analysis. The newest fully expanded leaf was placed in the measuring cuvette of a LiCor 6800 and photosynthesis assessed as a function of internal carbon dioxide concentration (*C*_i_) and as a function of light intensity. For full details, see supplementary methods.

Chlorophyll fluorescence assessment of *met1-1* mutants was carried out using a CFImaging system (Technologica Ltd, Colchester, UK). After 7 days of acclimation, four plants (one from each group) were analysed at a time and exposed to a light response curve protocol of decreasing light steps start at 1500 μmol m^-2^ s^-1^. For full method, see supplementary methods.

### DNA extraction- CTAB method

A modified version of the CTAB extraction protocol described by Porebski, Bailey and Baum (68) was utilized. For full details see supplementary methods.

### Whole Genome Bisulfite Sequencing

5µl genomic DNA from multiple samples per regime (n=6 plants/regime) were pooled together (total volume 30ul) to form a single sample. Library preparation and sequencing were carried out by Novogene.

Differentially methylated regions were computed with DMRcaller (69) using the bins method, with a bin size of 150 base pairs as done previously (13, 70). A p-value threshold of 0.01 for all contexts was used, alongside a minimum cytosine count of 4, minimum proportion difference of 0.2 for CpG and CpHpG, and 0.1 for CpHpH, and minimum reads per cytosine of 4, allowing bins with few cytosine bases to be avoided (71).

### RNA extraction and sequencing

The RNA from 3 independent replicates (100mg/replicate) per regime were extracted using the Macherey-Nagel Mini kit for RNA purification (Macherey-Nagel, Allentown, PA). Sample purity was assessed using the NanoDrop ND-1000 Spectrophotometer (NanoDrop, ThermoFischer, Wilmington, DE). Library preparation (poly-A) and sequencing were carried out by Novogene. Sequencing was performed on Illumina NovaSeq 6000 (Illumina, California, USA), utilizing a paired end 150bp read length, with 6Gb raw data generated per sample.

Raw data files were aligned to TAIR10 (72) using HISAT2 (73) and DESeq2 (74) was used to detect differentially expressed genes (DEGs) (p-adjusted value ≤0.05, and a log2-fold change ≥ 0.5).

## Supplementary Data

### Data and materials availability

All WGBS and RNA-seq data sets from this study have been submitted to the NCBI Gene Expression Omnibus (GEO; http://www.ncbi.nlm.nih.gov/geo/) under accession number GSE261533.

## Acknowledgments

We thank Dr. Marco Catoni for sharing of *met1-1* seeds and for comments on the manuscript. The analysis was performed on the HPC at the University of Essex, and we thank Stuart Newman for his support on using the cluster.

## Author contributions

R.A.E., T.L., U.B. and N.R.Z. conceived and designed the experiments. R.A.E. performed the experiments and analysed the data. T.L., U.B. and N.R.Z. supervised the work. R.A.E., T.L. and N.R.Z. wrote the paper. The authors read and approved the final manuscript.

## Supplementary methods

### Light Regimes

Growth light regimes utilised were the same as in Vialet-Chabrand et al (27). In short, the fluctuating light regime was recreated from recording natural light intensity during a relatively clear day in July 2011 at the University of Essex (Colchester, UK). This data was used to programme into a Heliospectra LED light source, representing the FLH regime. The average intensity of this regime was 460 µmol m−2 s−1, and this value was utilised as the intensity over the square high light regime (SQH). For the low light regime (FLL), the FLH regime was halved, representing an average of 230 µmol m^−2^ s^−1^ which was utilised for SQL.

### Chlorophyll Fluorescence

Four plants, one from each regime, were placed in the imager at a time. Measurements were all conducted between the 8am and 3pm. Plants were first dark adapted for at least 20 min prior to minimal fluorescence (*F_o_*) being measured by a weak measuring pulse. The maximal fluorescence (*F_m_*) was captured follow exposure to an 800 ms saturating pulse of 6231 μmol m^-2^ s^-1^. These measurements were used to determine the maximum quantum efficiency of PSII photochemistry with the following equation *F_v_/F_m_* = (*F_m_*- *F_o_*)/*F_m_*. This was followed by a light response curve, starting at PPFD value of 1500 μmol m^-2^ s^-1^, decreasing to 1250, 1000, 500, 250,150, and finally 50 μmol m^-2^ s^-1^. *F’* was constantly monitored, with *F_m_’* taken 3 minutes after each light level by applying a saturating pulse.These parameters were used to calculate *F_v_/F_m_, F_q_’/F_m_’, F_v_’/F_m_’, F_q_’/F_v_’*, and NPQ (75).

### Infrared Gas Exchange Analysis

For all gas exchange measurements, a Li-Cor 6800 portable gas exchange system (Li-Cor, Lincoln, Nebraska, USA) was utilised. A constant flow rate of 500 μmol m^-2^ s^-1^ was used, with constant cuvette conditions of 400 μmol mol^-1^ [CO_2_], vapour pressure deficit of 1.2 (±0.2) kPa and leaf temperature of 23°C unless otherwise stated. All measurements were taken between 8 am and 3 pm on the youngest fully expanded leaf on days 16-20 of exposure to light regimes. Differences in leaf area within the chamber were accounted for by determining the coverage of the chamber by photographing the leaf position in the chamber and analysed using open-source image analysis software ImageJ version 1.52a (76).

### A/Ci Response Curves

Assimilation rate (*A*) measured as a function of intercellular [CO_2_] (*Ci*) known as *A*/*Ci* curves were measured at 1500 μmol m^-2^ s^-1^ Photosynthetic Photon Flux Density (PPFD). Leaves were first stabilised in the chamber at ambient [CO_2_] of 400 μmol mol^-1^, and once stable, a measurement was taken, after which, [CO_2_] was decreased to 300, 200, 100, 80, and 50 μmol mol^-1^ before returning to ambient [CO_2_]. This was followed by an increase in [CO_2_] to 550, 700, 900, 1100, 1300, and 1500 μmol mol^-1^. Measurements were taken at each new [CO_2_] when *A* had stabilised which was usually within 30-120s.

### Light Response (A/Q) Curves

The response of *A* to changes in light intensity, was measured under the same conditions as *A/Ci* above, with the CO_2_ concentration maintained at 400 μmol mol^-1^ throughout the measurements. Leaves were stabilised at 1800 μmol m^-2^ s^-1^ to reduce stomatal limitation and a measurement recorded, and then PPFD was decreased to 1500, 1300, 1100, 900, 700, 550, 400, 250, 150,100, 50, and finally 0 μmol m^-2^ s^-1^, with a measurement taken at each step when *A* was stable and before *gs* responded to the new light level.

### DNA extraction- CTAB method

Tissue was harvested between midday and 1pm on day 18-21 of exposure to light regimes.100 mg of leaf tissue from a single plant (fresh weight) was ground in liquid nitrogen, to which 1ml CTAB buffer warmed to 55°C was added. This was then incubated at 55°C for 15 minutes. 1 volume of chloroform was added, and samples were incubated on ice for 10 minutes. These were vortexed for 1 minute then spun at 14000 r.p.m. for 5 minutes, and the supernatant recovered, at which point 5ul RNAse A (10mg/ml) was added and incubated at 37°C for 15 minutes. 1 volume chloroform was added, mixed by inversion for 5 minutes, centrifuged at 14000 r.p.m. for 10 minutes, and the supernatant recovered. To this 2.5 volumes 70% ethanol and 0.1 volumes 3M Sodium Acetate were added, and samples incubated at −20°C for 1 hour. Samples were then spun at 14000 r.p.m. for 5 minutes, the supernatant discarded and resulting pellet washed with 70% ethanol. Once dried, the pellet was resuspended in pure water. DNA integrity and purity was assessed using NanoDrop ND-1000 UV-Vis spectrophotometer was used along with the NanoDrop V3.1 software (Labtech International Limited, UK).

### Whole genome bisulfiute sequencing

Genomic DNA was fragmented into 200-400bps using a S220 focussed-ultrasonicator (Covaris, Massachusetts, USA) and Bisulfite treated with EZ DNA Methylation Gold Kit (Zymo Research, California, USA). Library concentration was quantified by Qubit 2.0 Fluorometer (Invitrogen, ThermoFisher Scientific, Massachusetts, USA), diluted to 1ng/µl and insert size checked on Agilent 2100 before further quantification by qPCR. Libraries were then pooled and fed into NovaSeq 6000 (Illumina, California, USA), utilizing a paired end 150bp read length and 30x coverage sequencing depth, with 8Gb raw data generated per sample.

### WGBS and RNA-seq analysis

Raw data was processed through Trimmomatic (77) to remove adapter specific sequences, remove low quality reads, and reads below a minimum length. The resulting files were aligned with Bismark (78) and Bowtie2 (79), to Arabidopsis thaliana (Col-0) reference genome build TAIR10 (72). We then removed duplicates with Bismark deduplication method.

Functional annotation utilized TAIR10 (72) annotation. Several genomic functional annotations were utilized; introns, exons, 5’UTR, 3’UTR, transposable elements, promoters (1 Kb upstream of TSS), and intergenic regions encompassing regions outside of these annotations.

For DEGs, heatmaps were produced using pheatmap (80) and volcano plots using EnhancedVolcano (81).

## Supplementary Fig. Legends

**Fig. S1:**
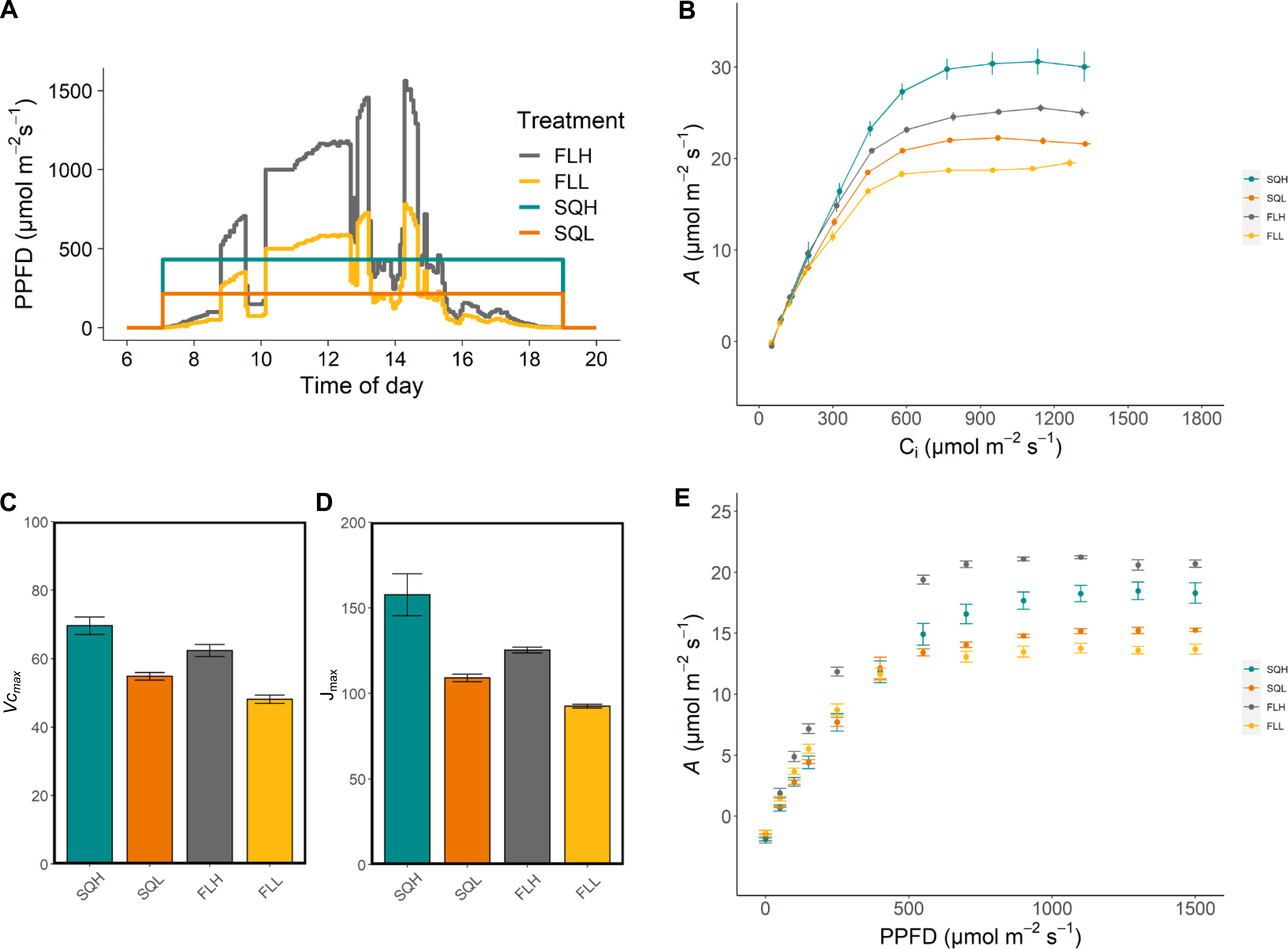
Diurnal light regimes utilised in this study alongside physiological assessment. *(A)* Area of the curves are equal, demonstrating the same average amount of light energy over a 12-hour period in low (square wave, SQL; fluctuating wave, FLL) and high light conditions (square wave, SQH; fluctuating wave, FLH). *(B)* Assimilation as a function of internal CO­_2_ concentrations in mature plants acclimated to each of the light regimes. From this, the maximum rate of carboxylation of Rubisco *(V­_cmax_; C)* which was significantly different across regimes, with both high light regimes having a greater rate than under low light, and fluctuating regimes performing worse than their square light counterpart (*p<0.05*). The maximum electron transport rate for RuBP regeneration *(J_max_; D)* was also calculated, which demonstrated a significant decrease in RuBP regeneration in low light acclimated plants compared to high light plants (*p<0.05*). *(E)* The effect of changing Photosynthetic Photon Flux Density (PPFD; μmol m^-2^ s^-1^) on photosynthesis in plants grown under the described light regimes. Letters on B and C indicate the results of Tukey post-hoc testing. Data shows the means ± SE (n=6 plants)

**Fig. S2:**
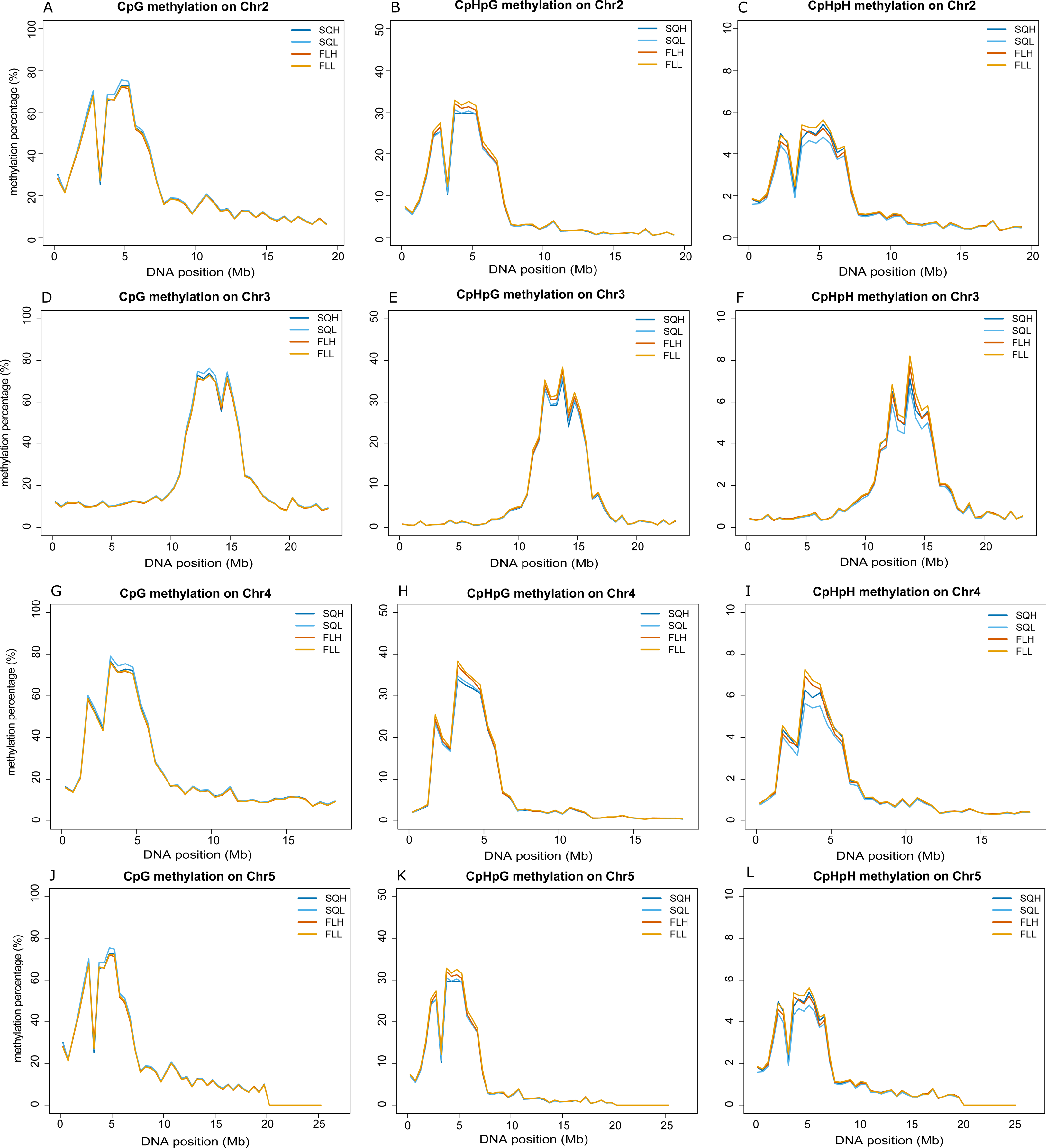
The effects of light regime on DNA methylation. Low resolution profiles of DNA methylation in the four light conditions. We considered separately CG, CHG and CHH methylation patterns and plot methylation data on chromosomes 2 to 5.

**Fig. S3:**
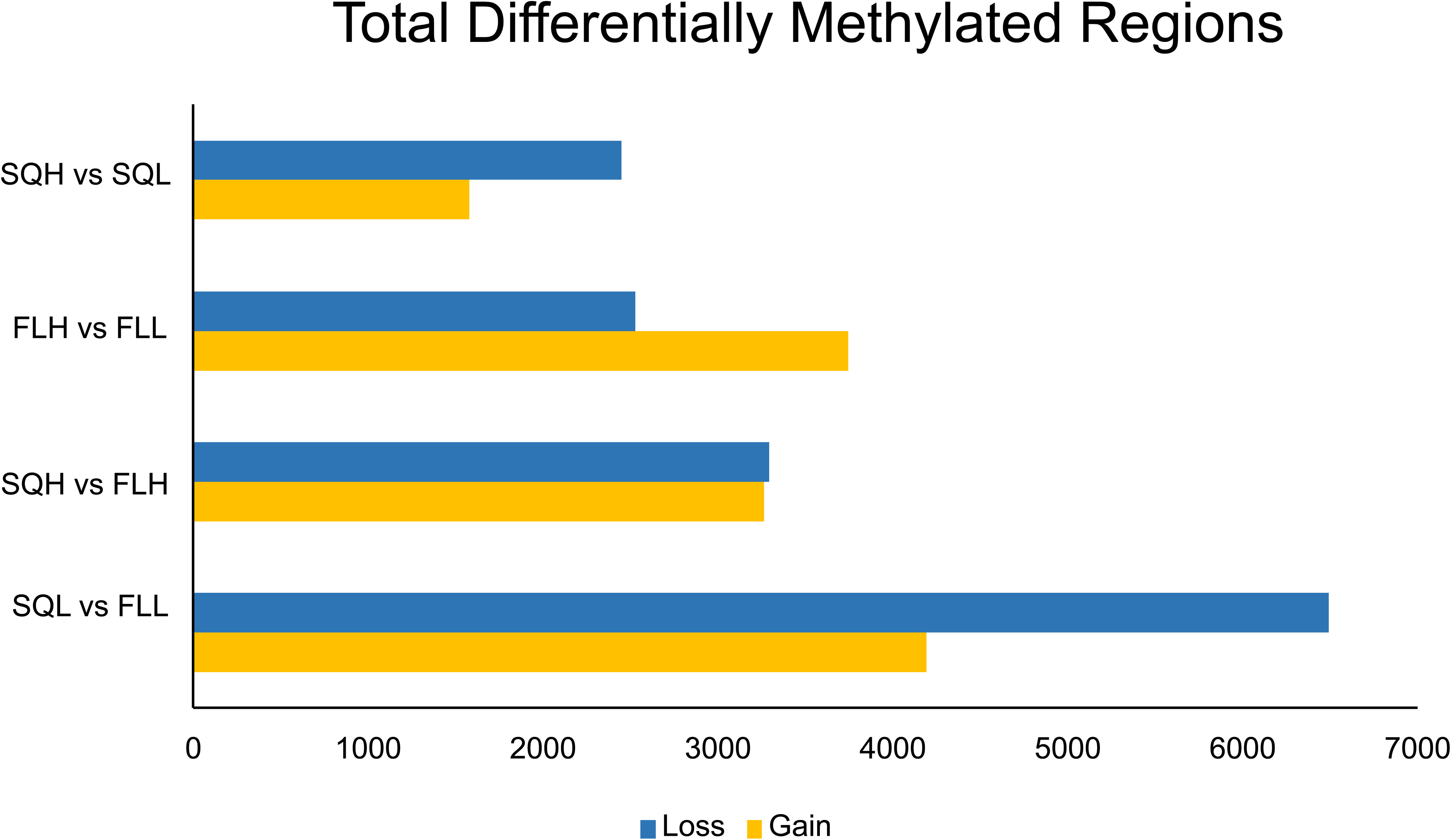
Number of DMRs in different comparisons. Barplot represents the number of DMRs that loss (blue) or gained methylation in the second light regime compared to the first one. We considered the comparison of light intensity exposure (SQH vs SQL and FLH vs FLL) and of light frequency (SQH vs FLH and SQL vs FLL). The barplot includes all DMRs in CpG, CpHpG and CpHpH contexts.

**Fig. S4:**
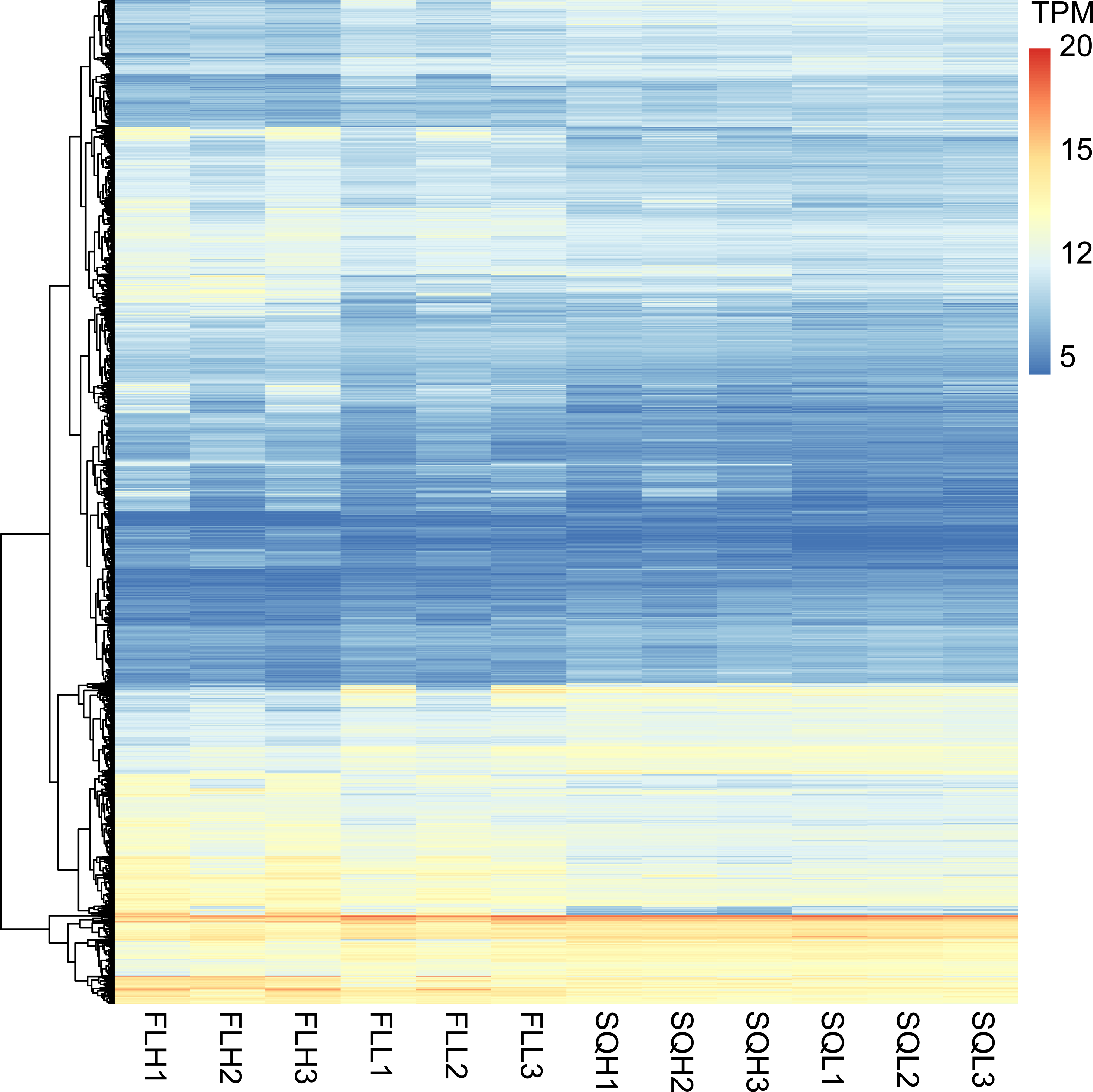
RNA-seq data in different light regimes. The heatmap shows transcripts per million (TPM) for each gene for each of the three replicates of the four light regimes.

**Fig. S5:**
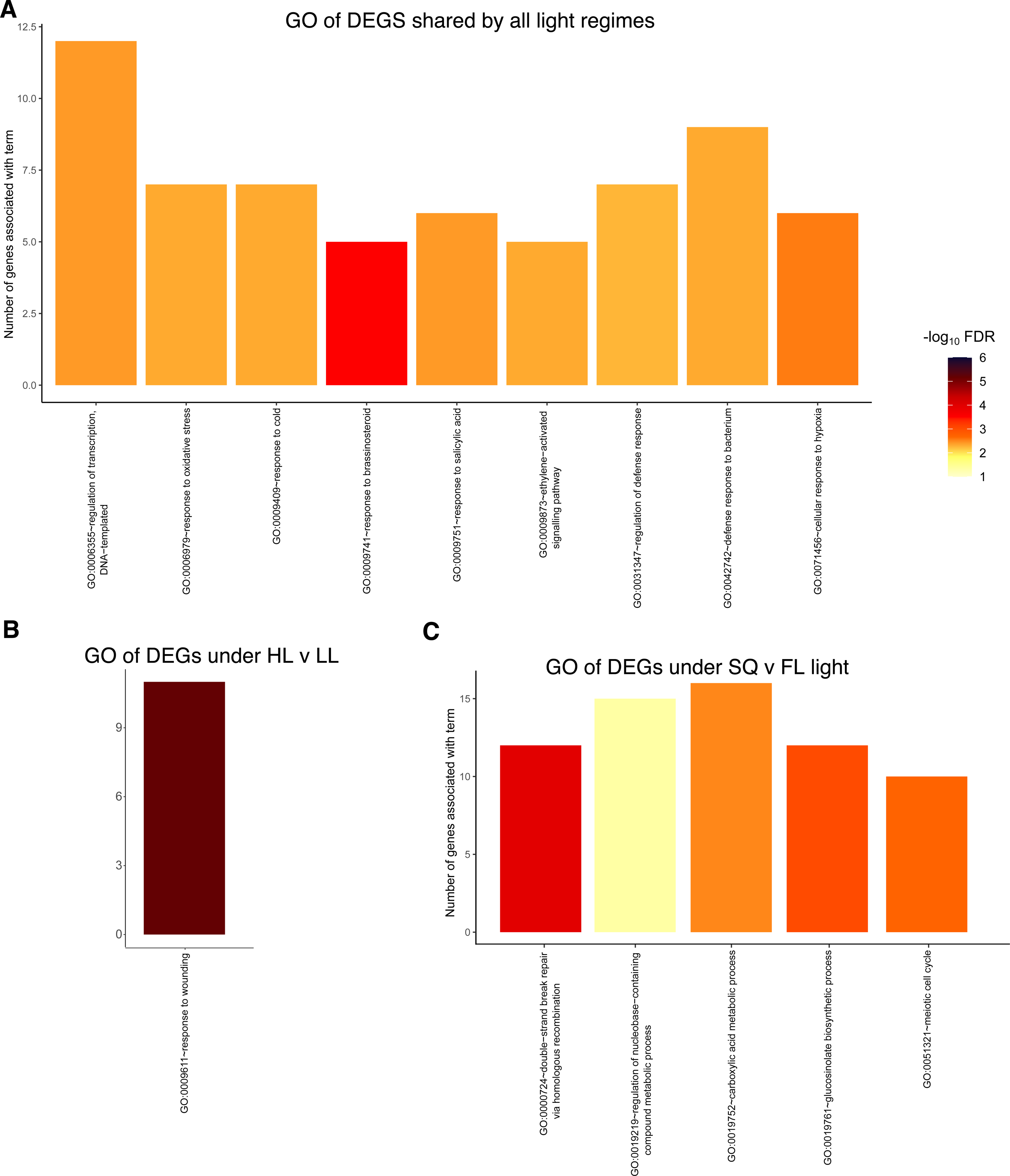
Gene ontology of overlapping differentially expressed genes between light regime comparisons. Bar height shows the number of genes associated with the indicated term, while colour indicated the -log_10_ of the False Discovery Rate (FDR).

**Fig. S6:**
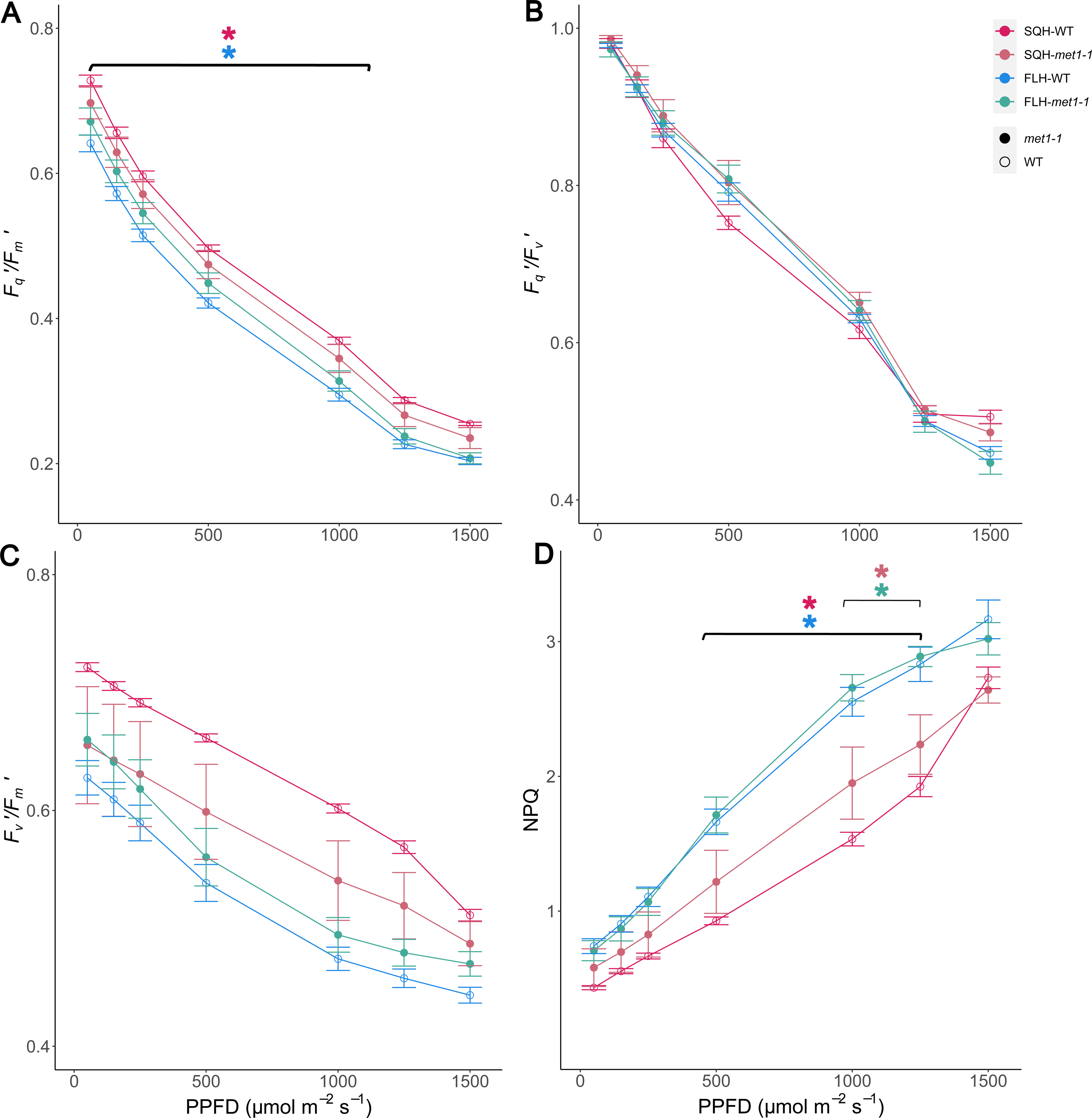
Chlorophyll fluorescence imaging of met1-1 and wild type Arabidopsis acclimated to SQH and FLH in response to changing light. *(A)* The operating efficiency of photosystem II (*F_q_’/F_m_’) (B)* Photochemical quenching (*F_q_’/F_v_’) (C)* Maximum efficiency of photosystem II (*F_v_’/F_m_’) (D)* Non photochemical quenching (NPQ). N=6, points represent the mean ± SE

## Supplementary Tables

**Table S1:** *List of light constitutive genes and TEs.* The 122 genes and TEs that harbour DMRs between all light regimes.

**Table S2:** *Functions of genes that are both differential methylated and differentially expressed in growth light regime comparisons.* The gene code, gene name, and function were obtained from TAIR. Context refers to the methylated cytosine(s) that changed in methylation between regimes.

**Table S3:** *List of differentially methylated and differentially expressed transposable elements.* Data is grouped by growth light regime and shows the cytosine contexts that display differential methylation, whether methylation is lost or gained, the log_2_ fold change of RNA-seq, and the associated gene function as described on TAIR.

**Table S4:** *Differentially expressed genes within 1Kb of a differentially methylated transposable element (TE)*. Data is grouped by growth light regime and shows the cytosine contexts that display differential methylation, whether methylation is lost or gained, the log_2_ fold change of RNAseq, and the associated gene function as described on TAIR.

